# RNA switch model for localization and translation of the myelin basic protein mRNA

**DOI:** 10.1101/2025.11.19.689361

**Authors:** Ved V. Topkar, Vivian Wu, Lana T. Ho, Nicholas Ambiel, Alex Valenzuela, Kailee Yoshimura, J. Bradley Zuchero, Meng-meng Fu, Rhiju Das

## Abstract

Oligodendrocytes myelinate the central nervous system by extending cellular projections that ensheath axons and elongate to form lipid-rich myelin. Classic studies visualizing RNA dynamics showed that myelin basic protein (MBP), one of the most abundant myelin proteins, is locally synthesized at the myelin sheath through the transport and local translation of *Mbp* mRNA. *Mbp* transport requires its 1.5-kb 3’ untranslated region (3’ UTR) and prior work identified candidate sub-sequences that may act as *cis*-acting transport stimulating RNA elements, including one with putative secondary structure. Here, a high-throughput reporter assay, dimethyl sulfate (DMS)-based RNA structure probing, and microscopy in primary rat oligodendrocytes identify a structured 127-nt region that we name the *Mbp* localization signal (MLS) as both necessary and sufficient for RNA enrichment to oligodendrocyte projections. Lysate pulldown experiments further identify hnRNP-F – a known constituent of the *Mbp* RNA granule that can suppress mRNA translation – as associated with the MLS; paradoxically, binding of this protein should compete with the ordered MLS RNA structure. These results suggest a model in which the MLS switches between two RNA conformations with distinct protein partners during the transition from *Mbp* mRNA transport to *Mbp* translation at the myelin sheath. Such regulation of RNA behavior by structure switching may generalize to other eukaryotic mRNAs whose behaviors shift across space and time.

**Significance Statement:** In the brain, oligodendrocyte cells generate myelin, a type of insulation that wraps around neuronal axons in order to facilitate fast electrical signaling. A critical step in myelination is the local translation of MBP (myelin basic protein) in the myelin sheath. This requires the transport of Mbp mRNA, an incompletely understood phenomenon that we revisit using two recent approaches for mRNA structure and function. We refine a 127-nt region that is necessary and sufficient for mRNA transport to the myelin sheath. A proteomic screen reveals that this myelin localization signal (MLS) associates with a translation-suppressing protein called hnRNP-F, suggesting a model where Mbp mRNA switches between two states, one for transport and one for translation at the myelin sheath.

## Introduction

Myelination is a powerful mechanism that evolved in jawed vertebrates to facilitate fast conduction of electrical signals along axons without the need to increase axon diameter. The biogenesis of myelin in the central nervous system by oligodendrocytes involves the extension of projections from the cell body, the ensheathment of neuronal axons by these projections, and the subsequent concentric wrapping of many layers of membranes around the axon. Within the myelin sheath, microtubules facilitate the transport of many cargos, including organelles, proteins, and mRNAs (1–3). The delivery of these cargos is essential for building the myelin sheath, a biochemically complex cellular compartment that is enriched in specialized lipids and proteins.

Myelin Basic Protein (MBP) is the second most abundant protein in the myelin sheath (4) and *Mbp* is the most abundant mRNA expressed by oligodendrocytes (5). As a polybasic protein, MBP performs the critical function of myelin compaction, the process by which cytoplasm is extruded from the myelin sheath in order to form a more efficient insulator (6). Indeed, spontaneous deletion of *Mbp* in mice causes devastating defects in myelin compaction leading to a shivering pathology and lethality a few weeks after birth (7, 8).

Early myelin fractionation experiments indicated that MBP is a locally translated protein (9). Later cell culture studies established that *Mbp* mRNA indeed localizes to oligodendrocyte projections (10), that this occurs through active transport on microtubules (11–13), and that *Mbp* mRNA localized to oligodendrocyte projections is translated by colocalized translational machinery (14). This research into *Mbp* provided one of the first examples of RNA transport and localization, but the molecular mechanisms underlying these phenomena remain mysterious.

*Mbp* mRNA contains a short ∼150-nt 5’ UTR followed by a ∼450-nt coding region, flanked by a ∼1500-nt 3’ UTR. Early molecular dissection identified the 3’ UTR as necessary and sufficient for RNA localization to oligodendrocyte projections. This research proposed sub-sequences in the *Mbp* 3’ UTR that may be responsible for the 3’ UTR’s ability to stimulate RNA localization to oligodendrocyte projections, including an RNA Transport Signal (RTS) sufficient for transport into processes by binding hnRNP A2 (heterogeneous nuclear ribonucleoprotein A2) but insufficient for distal localization of the transcript (15). A separately proposed RNA Localization Region (RLR) was found to be necessary for distal localization, but was not tested for sufficiency. RLR alignments across species have not revealed conserved sequences motifs, but were noted to fold into similar structures in silico across human, mouse, and rat sequences, leading to the still untested suggestion that an RNA secondary structure in this region might be important for *Mbp* mRNA localization (15).

Here, we used dimethyl sulfate (DMS) mutational profiling with sequencing (DMS-MaPseq) and a massively parallel reporter assay to map the secondary structure of the *Mbp* 3’UTR and identified a structured sub-sequence as a minimal RNA localization signal. Using a reporter assay read out by single molecule RNA microscopy, we demonstrate that this *Mbp* localization structure (MLS) is necessary and partially sufficient for the RNA localization effects of the full-length *Mbp* 3’UTR. Pulldown-proteomics reveals that the MLS binds hnRNP-F, a known constituent of the *Mbp* RNA transport granule involved in translational repression. We conclude with a proposed model of how hnRNP-F interaction with the MLS can function as a switch between *Mbp* transport and local translation during myelination.

## Results

### The 127-nt MLS region is required for *Mbp* mRNA transport

We first mapped the native secondary structure of the entire *Mbp* 3’ UTR using dimethyl sulfate (DMS) mutational profiling with sequencing (DMS-MaPseq) (16) applied to primary *Rattus norvegicus* oligodendrocytes (Fig. 1A). Through high throughput sequencing and mutational profiling of DMS modification sites, we obtained a normalized per-nucleotide reactivity for all As and Cs along the entire *Mbp* 3’ UTR in primary oligodendrocytes (Fig. 1B, top) which we used to generate a data-informed secondary structure model (Fig. 1C).

**Figure 1:**
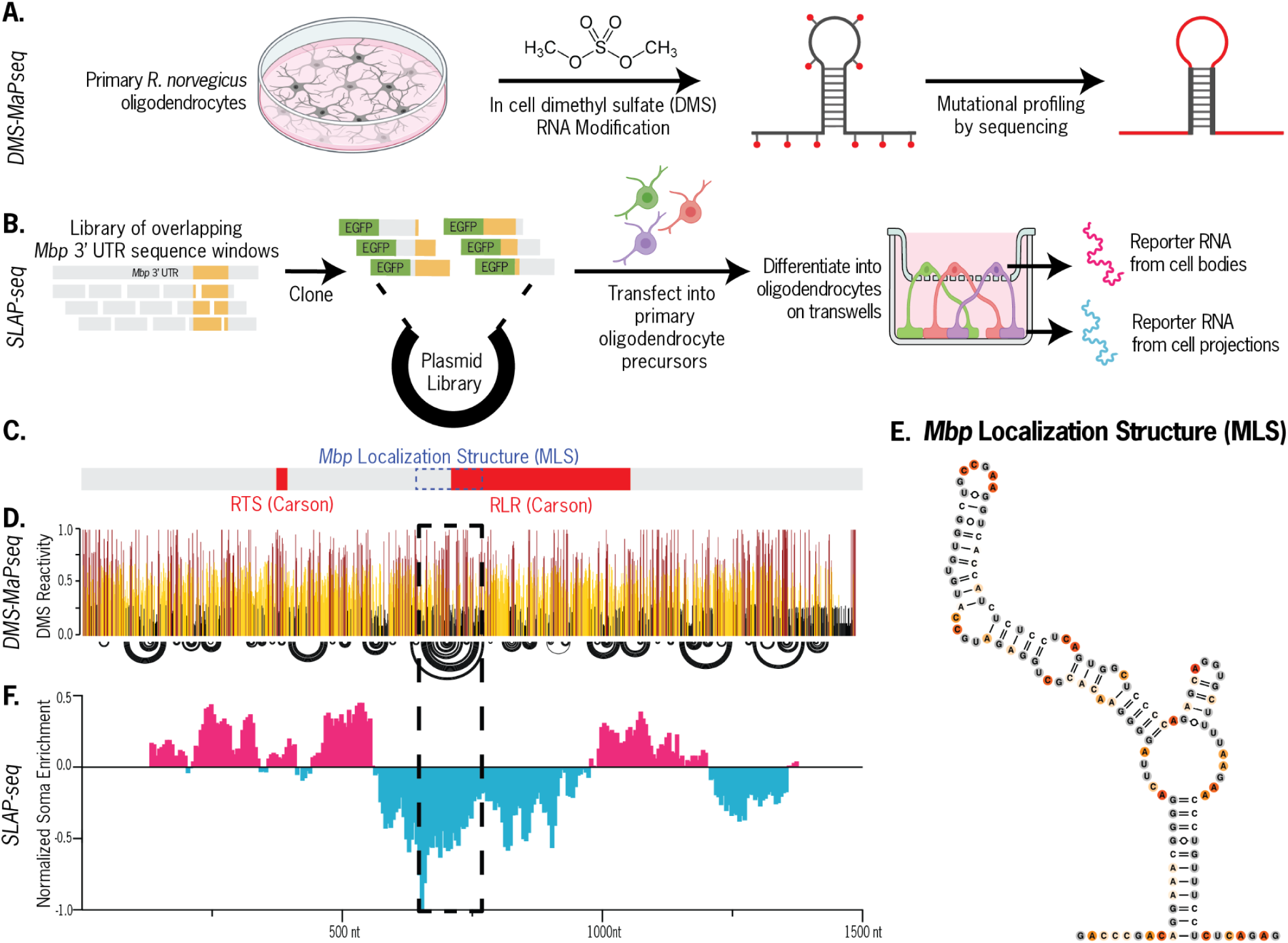
The *Mbp* localization signal (MLS). Schematics of (A) DMS-MaPseq protocol for mapping the secondary structure of the Mbp 3’ UTR in primary oligodendrocytes and (B) SLAP-seq to test a library of 3’ UTR sequences in their ability to stimulate reporter enrichment to oligodendrocyte processes. (C) RNA Transport Sequence (RTS), RNA Localization Region (RLR) and MLS locations along Mbp 3’ UTR. (D) In-cell DMS reactivity (top) and reactivity-guided secondary structure (bottom). Arcs represent base pairing between distal nucleotides. (E) Secondary structure of the MLS subsequence. (F). SLAP-seq reporter enrichment of windows along the Mbp 3’ UTR in cell bodies vs. projections. (G)Representative micrographs from reporter assay testing MLS role in peripheral RNA enrichment (scale bar = 30 μm), and (H) quantitative analysis supporting MLS necessity and sufficiency.

We separately sought to identify sub-sequences in the *Mbp* 3’ UTR that are functional in stimulating RNA localization, and developed <u>S</u>elective <u>L</u>ocalization <u>A</u>ssessed by <u>P</u>rojection <u>seq</u>uencing (SLAP-seq). We transfected primary rat oligodendrocyte precursor cells with a library of ∼270 reporters encoding *Egfp* followed by overlapping 250-nucleotide windows of the *Mbp* 3’ UTR, then differentiated them on 1-µm transwell filters. We then isolated bulk RNA from cell bodies and projections and sequenced them separately to calculate the relative enrichment of every 3’-UTR window between the two cell compartments (Fig. 1B). As expected, the previously identified RTS was insufficient for *Mbp* mRNA enrichment in oligodendrocyte projections (Fig. 1E). However, we identified multiple 3’ UTR sub-sequences sufficient to enrich an RNA to oligodendrocyte projections. Of note, regions 520-770 nt and 525-775 nt demonstrated sufficiency for distal localization to projections, though overlapping fragments 400-650 nt and 405-655 nt did not (Dataset S1). By comparing these two datasets, we identified the *Mbp* Localization Signal (MLS), a 127-nt region of the *Mbp* 3’ UTR (650-776 nt) that forms a well defined secondary structure in oligodendrocytes and represents the strongest sub-sequence of the full-length *Mbp* 3’ UTR in its ability to act independently as a *cis*-acting RNA localization signal. Of note, the MLS is a subset of the previously hypothesized RLR signal.

We validated that the MLS is necessary and sufficient for distal localization using a reporter assay in primary oligodendrocytes read out by single molecule fluorescence in situ hybridization (smFISH) (Fig. 2A and B).

**Figure 2.**
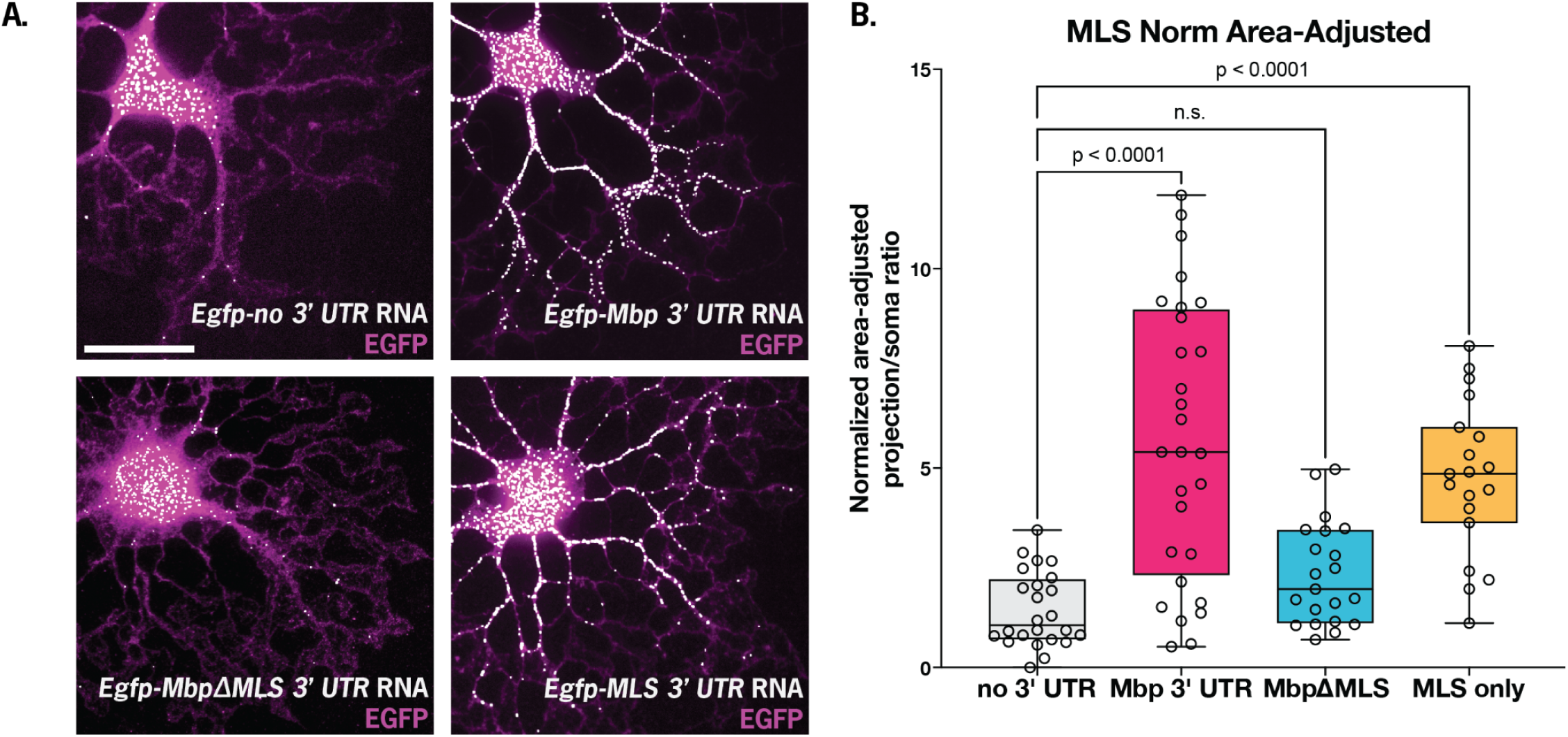
MLS is necessary and sufficient for distal localization of an EGFP reporter in primary oligodendrocytes. (A) Representative micrographs from reporter assay testing MLS role in peripheral RNA enrichment (scale bar = 30 μm), and (B) quantitative analysis supporting MLS necessity and sufficiency.

### Proteomic screen of MLS-associated proteins

We next asked how the MLS might function as a cis-acting RNA localization signal and set out to identify any potential MLS-binding proteins. We used a pulldown-proteomics approach by mixing biotinylated MLS RNA with cleared primary oligodendrocyte lysate and then pulling down on MLS-binding proteins via biotin binding with streptavidin immobilized on agarose beads. We then identified MLS-bound proteins using liquid chromatography-mass spectrometry (Fig. 3). When comparing proteins bound to the MLS vs. a scrambled negative control, the known mRNA-binding protein hnRNP-F was among the most enriched (Fig. 3, log_2_FC 1.48, p_adj_ < 0.002, Dataset S2). Though earlier work identified hnRNP-F as a protein present in *Mbp* transport granules (17), to our knowledge this is the first evidence of the protein associating with a specific region of the *Mbp* mRNA.

**Figure 3.**
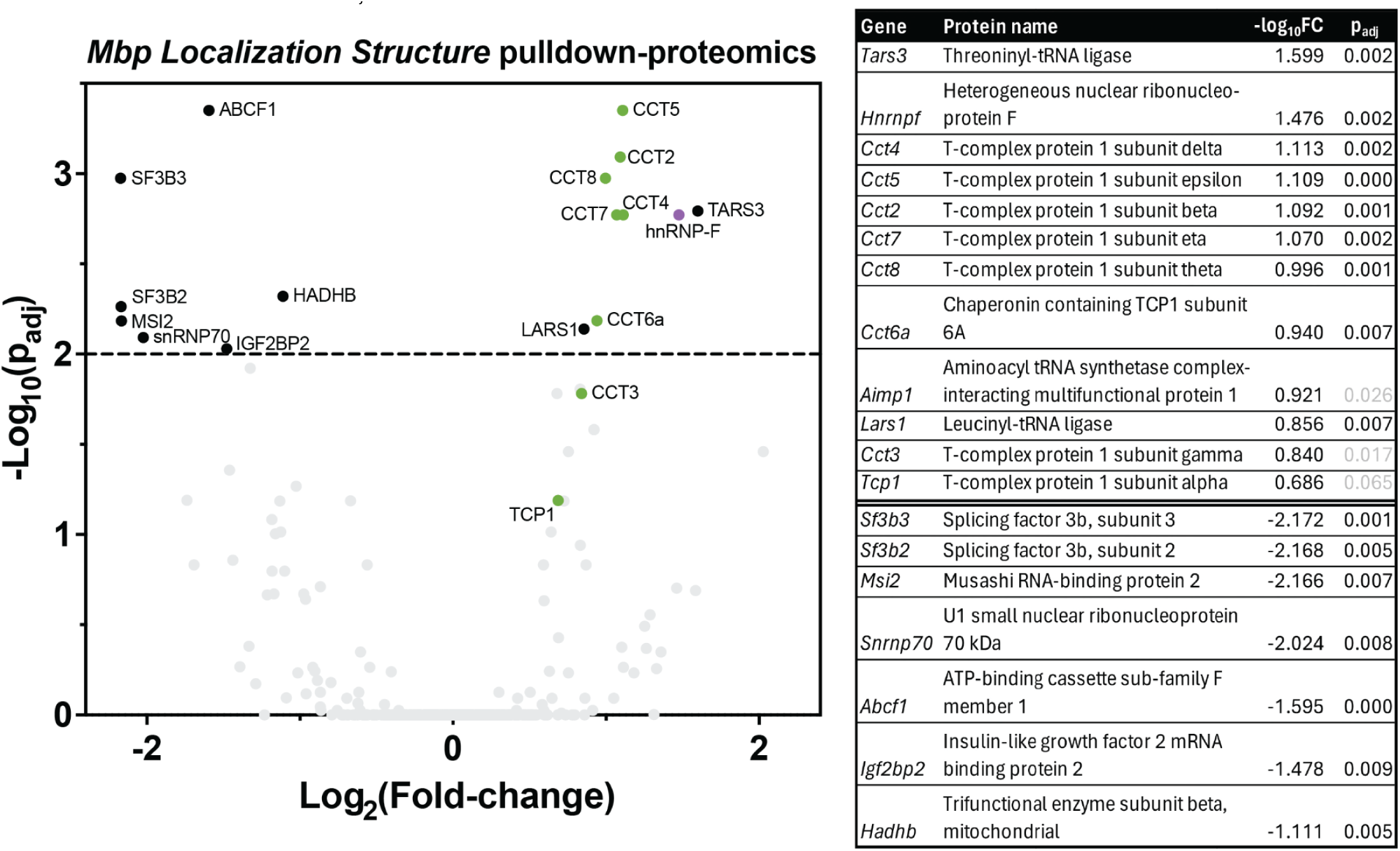
The MLS is a *cis*-acting inhibitor of RNA translation that interacts with hnRNP-F. (A) Volcano plot of mass spectrometry results of MLS RNA pulldown vs a scrambled negative control. hnRNP-F is significantly enriched at a threshold of p_adj_ < 0.01 (above dashed line). TRiC complex proteins (green) are also enriched.

In addition, we identified several components of the cytoplasmic protein translational apparatus and the TRiC protein folding chaperone complex, suggesting that the MLS may also be important for regulation of MBP translation. Other classic RNA-binding proteins including spliceosome-related proteins were significantly de-enriched, which may indicate functional avoidance of these proteins by the MLS or incidental sequence specificity of these proteins to the negative control bait.

### MLS and hnRNP-F interact to regulate *Mbp* translation

hnRNP-F is a known constituent of the *Mbp* RNA transport granule that inhibits MBP translation. Upon phosphorylation by Fyn kinase, it dissociates from *Mbp* granules (11), thereby activating local MBP translation (18). By immunostaining primary oligodendrocytes, we confirmed that hnRNP-F exists in a punctate pattern in both cell bodies and projections (Fig. 4A). To test the relationship between the MLS and translation suppression by hnRNP-F, we returned to our reporter assay and asked if MLS presence in the reporter 3’ UTR alters EGFP protein output. Indeed, the highest translational activity was for our negative control (no 3’ UTR). The addition of the MLS (alone or within the entire *Mbp* 3’ UTR) decreased the median translational output by ∼5-fold (Fig. 4B). Interestingly, adding the *Mbp* 3’ UTR without the MLS decreased median translational output by ∼2.5-fold, which suggests the presence of other, redundant cis-regulatory translation suppressors. Overall, we find that the MLS is sufficient and partially necessary for translational suppression by the *Mbp* 3’ UTR.

**Figure 4.**
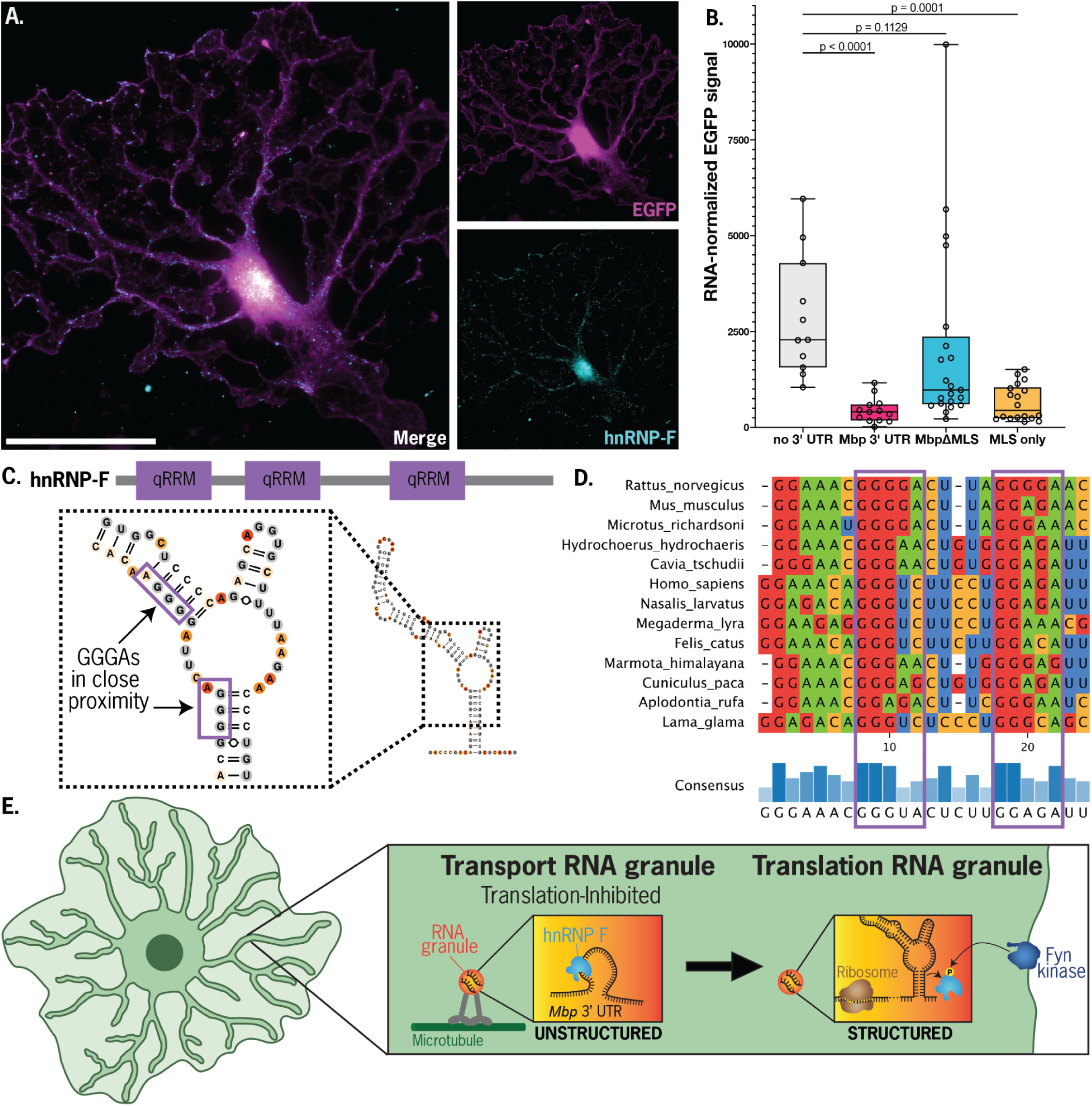
(A) Representative micrographs of an oligodendrocyte expressing cytoplasmic EGFP and stained with anti-hnRNP-F antibody (scale bar = 50 μm). In contrast to cytoplasmic EGFP, hnRNP-F is highly punctate, localizing both proximally and at the leading edge. (B)Testing MLS necessity and sufficiency in regulating reporter translation. Signal is the ratio of EGFP fluorescence to the number of reporter RNA copies per cell. Addition of the full-length Mbp 3’ UTR or the MLS to the 3’ end of an EGFP reporter dramatically reduces per-RNA translational output. Deletion of the MLS from the full-length Mbp 3’ UTR partially rescues translation, indicating that the MLS is sufficient for translational suppression, but not necessary. (C) Architecture of the hNRNP-F protein, which contains three qRRM RNA-binding domains with sequence specificity of 5’-GGG-3’. Enlarged MLS structure with GGG(A) sub-sequences highlighted. (D) Conservation of GGG(A) motif (purple boxes) in *Mbp* 3’UTR across mammalian species. (E) RNA switch model: during *Mbp* mRNA granule transport into oligodendrocyte projections, hnRNP-F binds to GGG(A) sequences in the unstructured *Mbp* 3’UTR, inhibiting translation. Once the granule localizes to the periphery of the cell, Fyn kinase phosphorylates hnRNP-F, resulting in its release from *Mbp* RNA and allowing the folding of the mRNA into the predicted translation-ready secondary structure.

The hnRNP-F protein architecture includes three qRRM domains that bind to single-stranded GGG RNA sequences with strongest affinity for GGGA (19). Indeed, the rat MLS contains two separate base-paired GGGA sequences in close proximity in the primary sequence, and most mammal *Mbp* MLS regions preserve one or both of these G-rich segments (Fig. 4D, Dataset S3). However, these are not single-stranded in the DMS-based secondary structure model (Fig. 4C), an apparent contradiction that we discuss below in our proposed model for hnRNP-F regulation of MBP translation.

## Discussion

Here, we developed a pipeline for studying mRNA structure and interactions during cellular localization using two orthogonal techniques. First, we applied biochemical RNA structure mapping to derive the secondary structure of the *Mbp* 3’ UTR in its native context of primary differentiating oligodendrocytes. We found a landscape of structured and unstructured regions in this 1.5-kb stretch of RNA and discovered that the secondary structure previously proposed in the literature diverges from our experimentally-informed structure. To our knowledge, this is the first application of the DMS-MaPseq method to primary mammalian cells. Second, we complemented this approach with a massively parallel reporter assay SLAP-seq, which confirmed that a small 127-nt region is indeed necessary for Mbp mRNA localization to oligodendrocyte projections.

Though the MLS is sufficient for *Mbp* mRNA localization (Fig. 2), we were surprised that our MLS-pulldown identified mainly proteins involved in translation (Fig. 3) but did not find motor proteins. This result suggests that the MLS does not directly bind with motor proteins, but may instead indirectly associate with motor proteins via intermediates like adapter proteins. For example, in neuronal dendrites, CPEs (cytoplasmic polyadenylation elements) sequences in mRNA bind to CPEB (CPE binding protein), an adapter protein that in turn associates with kinesins and dyneins. Interestingly, CPEB also associates with maskin, a protein that may regulate cap-dependent translational repression (20). Our findings here that hnRNP-F associates with the MLS and that the MLS alone reduces translation of an mRNA reporter, combined with the known association of hnRNP-F with motor proteins, is consistent with the prevailing dogma that transport mRNA granules are translationally repressed. Canonical examples include neuronal synaptic plasticity (21, 22) as well as RNA localization in *Drosophila* embryos (23, 24).

On first glance, the finding that the MLS forms a well-defined structure within oligodendrocytes appears to contradict with the proposed association of hnRNP-F with the MLS, which would need to expose GGG motifs that appear sequestered in the proposed RNA structure. To resolve this contradiction, we propose an “RNA switch model” for the *Mbp* RNA (Fig. 4E): the MLS element takes an unstructured form during *Mbp* biogenesis and transport, where it binds hnRNP-F and other factors that inhibit mRNA translation. When the *Mbp* mRNA transport granule reaches its destination at the cell periphery, membrane-bound Fyn kinase phosphorylates hnRNP-F, which dissociates from the *Mbp* mRNA and becomes translation competent (17, 18). The MLS, upon release of hnRNP-F, folds into a stable secondary structure that recruits translation factors. Although an RNA switch has not been previously proposed for the *Mbp* 3’ UTR, other protein-coupled RNA conformational changes have been identified across numerous processes involving non-coding RNA in mammalian cells, including splicing (25) and ribosome assembly (26). Future advances in ribonucleoprotein structure determination may enable the incisive testing and visualization of these multiple proposed states.

## Materials and Methods

Primary oligodendrocytes precursor cells were isolated through immunopanning from P5-7 rat cortices and differentiated *in vitro* for 3-5 days. Chemical mapping experiments were performed by treating cells with 1.6% dimethyl sulfate in 200 mM sodium bicine pH 8.5 at 37 C for 4 minutes then quenching with 30% 2-mercaptoethanol. Transfections were performed using the Amaxa Nucleofector kit for primary mammalian glial cells. RNA extractions were performed with Trizol LS (Thermo Fisher Scientific). Sequencing was performed on an Illumina Miseq. smFISH probes targeting the *Egfp* gene were purchased from Stellaris.

LC/MS was performed on an Orbitrap Exploris 480 mass spectrometer (Thermo Fisher Scientific). Data analysis was performed using Python, R, Fiji, Graphpad Prism, and RNAFramework. See details in SI Appendix.

### Primary cell isolation and culture

Primary oligodendrocyte precursor cells were isolated and cultured as described previously (27). Briefly, P5-P7 rat cortices were minced and enzymatically digested with papain under carbogen (95% O_2_ 5% CO_2_). The digestion was quenched with ovomucoid inhibitor and the resulting cell slurry was triturated and passed through a 20-µm Nitex mesh filter to remove cell clumps and isolate single cells. The cell suspension was then depleted of astrocytes and oligodendrocytes via immunopanning with anti-Ran2 and anti-GalC antibodies, respectively. Oligodendrocyte precursor cells were then isolated via immunopanning with anti-O4 antibody and released from immunopanning plates with trypsin (27).

OPCs were then cultured at 37 °C, 10% CO_2_, in DMEM-Sato growth medium (1X Dulbecco’s Modified Eagle’s Medium, 2 mM Glutamine, 100 U/mL penicillin, 100 µg/mL streptomycin, 1 mM Sodium pyruvate, 5 µg/mL insulin, 5 µg/mL N-acetyl-L-cysteine, 1X trace elements B, 10 ng/mL d-Biotin) supplemented with either proliferation (PDGF, NT-3, CNTF, forskolin) or differentiation (CNTF, forskolin, T3) growth factors.

For all imaging experiments, OPCs were grown in differentiating conditions on PDL-coated glass coverslips in 24-well plates at a density of 2,500 cells/coverslip. After 4 days of culture in differentiation conditions, the coverslips were fixed in 4% paraformaldehyde at room temperature for 10 minutes, then washed 3 times in PBS. Cells were permeabilized in 0.1% Surfact-AMP Triton-X in 1X PBS at room temperature for 10 minutes, then washed 3 times in PBS.

### DMS-MaPseq in primary oligodendrocytes

DMS modification was carried out using a sodium bicine buffering system base as previously reported (28). Primary rat OPCs were differentiated in 15-cm plates for 3-4 days, then modified by carefully replacing the media with a modification mixture (1X DMEM, 200 mM sodium bicine pH 8.5, 1.6% DMS). The plate was incubated at 37 °C for 4 minutes in a fume hood. Modification was terminated by removing the modification solution and quenching residual DMS with 3 washes of ice cold 30% BME v/v in PBS. Then, the cells were scraped down with a cell scraper into ice-cold PBS and pelleted at 1000 xg for 10 minutes at 4 °C. The pellet was resuspended in 1 mL TRIzol LS (Thermo Fisher) and cleaned up with the Direct-Zol RNA (Zymo Research).

For DMS-MaPseq library preparation targeted to the *Mbp* 3’ UTR, we first conducted reverse transcription under DMS mutational profiling conditions with TGIRT-III enzyme (29) using a reverse-transcription primer complementary to 3’ end of the the *Mbp* 3’ UTR. 1 µg of modified RNA was mixed with 10 pmol reverse transcription primer (oVT378) in water and denatured at 70 °C for 5 minutes and snap-cooled on ice.

Reverse transcription was then carried out in reactions containing 50 mM Tris-HCl pH 8.3, 75 mM KCl, 3 mM MgCl_2_, 1 mM dNTPs, 1 mM DTT, 100 U TGIRT-III enzyme (InGex) and incubated at 60 °C for 3 hours in a thermocycler with heated lid. Reactions were terminated and RNA hydrolyzed by addition of 200 mM NaOH, 1 mM EDTA and incubating at 65 °C for 15 minutes. The pH was neutralized by addition of Zymo Oligo Binding buffer and cDNA was then cleaned up with the Zymo Oligo Clean and Concentrator kit (Zymo Research) and eluted in water.

cDNA corresponding to the *Mbp* 3’ UTR was then PCR-amplified with NEBNext Ultra II Q5 Master Mix with oVT378 and oVT680 primers for 15 cycles and purified with RNACleanXP beads in a 1:1 ratio. The full 1.5-kb 3’ UTR dsDNA was gel purified with the Zymoclean Gel DNA Recovery Kits (Zymo Research). Library preparation for Illumina sequencing was carried out with the NEBNext Ultra II FS DNA Library Prep Kit using iTru ligation stubs and indexing primers (30). Briefly, 100 ng of dsDNA was enzymatically sheared using NEB FS Enzyme in a thermocycler with heated lid for 20 minutes at 37 °C, then inactivated at 65 °C for 30 minutes. iTru y-yoke ligation adapters containing overhangs for indexing PCR were denatured at 95 °C, then annealed by slow cooling at −0.1 °C/s to room temperature in a reaction containing 15 µM each iTrusR1-stub and iTrusR2-stub in 1X salty TLE (10 mM Tris pH 8.0, 0.1 mM EDTA, 100 mM NaCl). Sheared dsDNA was end-repaired and ligated to annealed iTru y-yoke ligation adapters by addition of 0.5 µM annealed iTru y-yoke, 1X FS ligation master mix + FS ligation enhancer at 20 °C for 1 hour. Ligated dsDNA was cleaned up with 0.9X AMPureXP beads. Indexing PCR was carried out with NEBNext Ultra II Q5 Master Mix for 5 cycles with 2 µM each of iTru5 and iTru7 indexing primers. This final sequencing library was cleaned up with 0.8X AMPureXP beads, quantified by Qubit (Thermo Scientific) and Bioanalyzer (Agilent Technologies) and sequenced on an Illumina MiSeq.

DMS-MaPseq analysis was done with the RNAFramework package (31). Sequencing reads were aligned to the *Rattus norvegicus* 3’ UTR sequence with bowtie2 (32) using the rf-map module, DMS modification-induced mutations were counted using the rf-count module, and then normalized with the rf-norm module using the Zubradt 2016 scoring method (29) with boxplot normalization, allowing each nucleobase to be normalized separately, and using a sliding normalization window of 50 nt. Finally, data-informed secondary structure prediction was carried out with the rf-fold module using RNAStructure as the prediction engine (33). G and U nucleotides were excluded due to poor signal-to-noise of DMS modification at these nucleobases in live cell treatment experiments.

### Selective Localization Assessed by Projection sequencing (SLAP-seq) for RNA localization

Random fragments of the *Mbp* 3’ UTR were generated by first amplifying a gene block of the *Mbp* 3’ UTR sequence (gVT9) for 15 cycles with oVT490/491 and agarose gel purifying the 1.5-kb product. 100 ng of dsDNA was enzymatically sheared using NEB Fragmentase Enzyme in 1X NEB Fragmentase Buffer in a thermocycler with heated lid for 20 minutes at 37 °C. Fragmentation was terminated by addition of 100 mM EDTA, then column purified with the Zymo DNA Clean and Concentrator 5 kit. The resulting dsDNA was end-repaired, dA tailed, and ligated to adapters containing restriction cloning sites with the Kapa HyperPrep kit. Ligation adapters were prepared by separately annealing 15 µM oVT493/494 (5’ adapter) or oVT495/496 (3’ adapter) in 1X salty TLE (10 mM Tris pH 8.0, 100 mM EDTA, 100 mM NaCl), first by denaturing at 95 °C for 2 minutes, then slowly cooling to 25 °C. End-repaired and dA tailed dsDNA was ligated to these annealed ligation adapters with the Kapa HyperPrep ligase enzyme at adapter:insert molar ratio of 100:1 at 20 °C for 4 hours. The ligation product was cleaned up with 1.0X AMPureXP beads, amplified for 15 cycles with oVT482/oVT537, and agarose gel purified to isolate DNA of size 200-400 bp. The dsDNA was then restriction digested with BsrGI and SacII enzymes, and purified with Zymo DNA Clean and Concentrator 5 columns.

The restriction digested inserts were ligated with ElectroLigase (New England Biosciences) into a double digested backbone plasmid with upstream EGFP ORF driven by the MBP promoter. The ligation contained a 3:1 insert:vector ratio and incubated at room temperature for 30 minutes before inactivation at 65 °C for 15 minutes. Ligated products were then electroporated into NEB 10-beta electrocompetent cells using a 2-mm cuvette with settings 2.0 kV, 200 Ohms, 25 µF and plated on 245-mm x 245-mm LB agarose plates. Roughly 200,000 colonies were collected and plasmid extracted with ZymoPURE Plasmid Maxiprep kit (Zymo Research).

For the transwell assay, transwell chambers were first prepared by first coating the bottom of 6-well plates with poly-D-lysine, then affixing 1-µm transwell filters in each well with clear nail polish. For each transwell chamber, 5 million proliferating OPCs were lifted with trypsin and nucleofector with the plasmid library with the Amaxa Nucleofector II using the Amaxa Basic Nucleofector Kit for Primary Mammalian Glial Cells. The entire nucleofection was plated on the top of the transwell chamber. This process was repeated for 6 total transwell chambers. The chambers were incubated in differentiation media with daily half feeds. After 4 days of differentiation, the top and bottom of each filter were rinsed with ice cold DPBS and the top of each filter was scraped with a micropipette tip to release lysate from the oligodendrocyte cell bodies. The transwell was then removed from the plate and the bottom of the well was scraped with a micropipette tip to release lysate from the oligodendrocyte projections. The compartment-specific lysate from 6 transwell chambers was combined. RNA from lysate was purified with the Direct-zol RNA Purification Kit (Zymo Research) with on-column DNase treatment.

Compartment-specific reporter RNA was library prepped with a protocol inspired by mcSCRB-seq (34). Reverse transcription with primers containing P7 adapter and i7 index sequence was carried out in reactions containing 10% PEG 8000, 1 mM dNTPs, 1 µM reverse transcription primer (oVT506-508), 1X Maxima RT Buffer, 20 U Maxima H Minus Reverse Transcriptase). Reverse transcription was incubated at 42 °C for 90 minutes, then cleaned up with RNACleanXP beads. The P5 adapter and i5 index sequence was added by limited cycle PCR in KAPA HiFi HotStart Readymix with 0.25 µM each oVT509-511 and iTru_P7 for 10 cycles. The PCR was cleaned up with 1X AMPureXP beads. These libraries were preliminarily quantified with qPCR and endpoint PCR was performed as needed with iTru_P5/iTru_P7 primers using NEBNext Ultra II Q5 Master mix. The final libraries were quantified with Qubit and Bioanalyzer and sequenced on an Illumina MiSeq sequencer. Separately, the input plasmid library was library prepped for sequencing by 2-step PCR amplifications. The first PCR was with oVT517/538 to add overhangs for addition of Illumina sequencing adapters. The second PCR was with iTru5/iTru7 indexing primers. The resulting sequencing library was quality controlled, quantified, and sequenced as above.

Demultiplexed reads were trimmed with CutAdapt (35), aligned against the *Mbp* 3’ UTR sequence with bowtie2 (32), and converted to BED files with BEDOPS (36). We used a rolling window approach to quantify the extent to which each sub-sequence of the *Mbp* 3’ UTR was enriched in cell projections. We defined overlapping windows of size 250 nt every 5 nt along the entirety of the 3’ UTR sequence. Then, the number of reads in each sample that entirely fall into every window was counted. Per-window counts for the projection or cell body fractions were divided by the per-window counts derived from the input plasmid library to obtain normalized per-window counts. The normalized enrichment was defined as the per-window difference between the normalized top and normalized bottom counts scaled such that the maximum value was 1.

### Sequence alignment

The determined *Mbp* localization signal (MLS) region, alongside previously-characterized projection-localizing sequences (10), was aligned to the rat *Mbp* 3’UTR NCBI Reference Sequence: NM_001025294.1 using SnapGene.

### smFISH staining and microscopy of endogenous transcripts

smFISH DNA probes were designed using thermodynamic parameters described previously (37) with FLAP-X overhangs, ordered in plate format, and pooled in equimolar concentration. To prepare pre-hybridized smFISH probes for cell staining, a 10-µL reaction of 40 pmol of probe pool, 50 pmol Cy3-labeled FLAP-X ssDNA, in 1X NEBuffer3 was mixed and hybridized in a thermocycler with heated lid by heating to 85 °C then slowly ramping down to 25 °C.

Cells were cultured, fixed, and permeabilized as above. To stain cells for a given RNA, coverslips were first incubated in wash buffer (10% formamide in 2X SSC) at room temperature for 15 minutes. A 50-µL hybridization mixture containing 48.5 µL hybridization buffer (2X SSC, 10% formamide, 10% w/v dextran sulfate, 0.2 µg/µL RNase-free BSA, 1 µg/µL *E. coli* tRNA, 2 mM Ribonucleoside Vanadyl Complex), 1 µL pre-hybridized smFISH probe mixture, and 0.5 µL of primary antibodies was pipetted on Parafilm in a glass container humidified with a wet paper towel. The coverslip was then carefully laid face-down on the hybridization mixture. The glass container was then sealed with Saran wrap incubated at 37 °C overnight in the dark. On the second day, the coverslips were transferred face-up in a 24-well plate and washed 3 times for 10 minutes each at 37 °C with 1X wash buffer. For secondary antibody and DAPI staining, the coverslip was laid face-down on a 50 µL mixture of hybridization buffer, secondary antibodies, and DAPI at final concentration of 300 nM and incubated at room temperature for 30 minutes. The coverslips were transferred back to a 24-well plate and washed 3 times for 5 minutes each at room temperature in 2X SSC. Coverslips were then mounted on glass slides with 5 µL of Prolong Glass mounting media and cured in the dark for at least 24 hours before imaging.

Slides were imaged with an 400X oil immersion objective on an Olympus IX83 epifluorescence microscope equipped with an Orca Fusion sCMOS camera. RNA spot detection was performed using the FISH-quant v2 package (38).

For representative images in main-text figures, the RNA channel was background-subtracted, denoised, binarized, and dilated in Fiji (39) for ease of visualization. All formal image analysis was performed on raw micrographs.

### Reporter construct cloning and imaging

Reporter plasmids encoded, in order, the UbC promoter, the *Egfp* ORF, a 3’ UTR sequence of interest, and finally the polyadenylation signal from the rat *Mbp* gene. To minimize toxicity caused by reporter overexpression as we observed when using a CMV or *MBP* promoter, we opted for the weaker UbC promoter. To maintain physiologic transcription, we used the *Mbp* polyadenylation signal. An initial plasmid encoding the *Mbp* 3’ UTR (pVT70) was constructed with Gibson cloning, and all subsequent plasmids were created by replacing the *Mbp* 3’ UTR with other 3’ UTRs using restriction cloning at the BsrGI and BamHI sites.

Transfection of reporter constructs was done with the Amaxa Nucleofector 4D. For each nucleofection, 4×10^5^ proliferating OPCs were nucleofected with 400 ng plasmid, then plated on poly-D-lysine coated coverslips at a density of 1×10^4^ cells/coverslip in differentiation media. Coverslips were fixed at culture day 4.

Fixed coverslips were stained as above, with the exception of using premade smFISH probes against the *Egfp* ORF as purchased from Stellaris. To better visualize EGFP expression in case of attenuation by harsh denaturing steps in the smFISH protocol, EGFP protein was stained with goat anti-GFP primary antibody (Rockland #600-101-215). hnRNP-F was stained with ProteinTech #67701. Tubulin was stained with Sigma #T9026. Slides were imaged with 100X oil immersion objective (NA 1.45) on an Olympus IX83 epifluorescence microscope equipped with an Orca Fusion sCMOS camera. RNA spot detection was performed as above. The cell mask was created by separately segmenting the cell on the Tubulin and EGFP channels using the Fiji Threshold tool, then merged together. Cell body segmentation was performed manually using the tubulin channel. All manual annotation was performed blinded to reporter construct identity and unblinding only occurred after final analysis was completed. To quantify RNA enrichment, a raw enrichment score was first computed by dividing the RNA puncta count outside vs inside the cell body. To adjust for differences in cell morphology between conditions caused by reporter expression, we divided the raw puncta enrichment by the ratio of cell projection area to cell body area. Finally, we normalized all area-adjusted ratios to set the median of the negative control (no 3’ UTR) to 1.

### RNA-Protein Pull-down Proteomics

Biotinylated RNA baits were prepared through *in vitro* transcription followed by enzymatic biotin labeling. DNA templates were designed with the T7 promoter followed by either the MLS sequence or a scrambled negative control and assembled by PCR assembly (40). 1 µg of each DNA template was *in vitro* transcribed with the TranscriptAid T7 High Yield transcription kit (Thermo Scientific) and cleaned up with Zymo RNA Clean and Concentrator 25 spin kit (Zymo Research). The RNA was then size-selected on a 12% denaturing urea PAGE gel and purified with the ZR small-RNA PAGE Recovery Kit (Zymo Research). Enzymatic biotin labeling was carried out using the Pierce RNA 3’ End Biotinylation Kit (Thermo Scientific). Each labeling reaction contained 500 pmol RNA and was allowed to incubate at 16 °C overnight. Labeling reactions were extracted with phenol/chloroform/isoamyl alcohol followed by ethanol precipitation.

Cell pellets were prepared by mechanically scraping down in 3-5 million primary rat oligodendrocyte precursor cells differentiated in vitro for 5 days into ice-cold D-PBS. The lysate was pelleted at 500 xg for 30 minutes at 4 °C. Pellets were lysed in lysis buffer (140 mM KOAc, 20 mM HEPES pH 7.4, 10 mM MgOAc, 1X HALT EDTA-free protease inhibitor cocktail, 1 mM DTT, 1% NP-40) at 4 °C for 30 minutes with gentle rocking, then cleared with centrifugation at 20,000 xg at 4°C for 30 minutes. Protein concentration was measured with the Pierce BCA Protein Assay Kit (Thermo Scientific). Pellets were collected in biological triplicates for each RNA bait.

Biotinylated RNA baits were denatured at 80 °C for 4 minutes, then snap-cooled on ice. RNA-protein binding reactions contained 100 pmol denatured RNA, 150 µg of cleared lysate, and 1X Binding/Wash buffer (140 mM KOAc, 20 mM HEPES pH 7.4, 10 mM MgOAc, 1X HALT EDTA-free protease inhibitor cocktail, 1 U/µL SUPERase-In RNase inhibitor, 1 mM DTT). Binding was carried out at room temperature for 1 hour with gentle rocking.

RNA-protein complexes were pulled down with Pierce High Capacity Streptavidin Agarose beads. Beads were first prepared by pelleting at 500 xg for 30 seconds and washed twice in 1X Binding/Wash buffer. Then, 50 µL of washed beads were added directly to RNA-protein binding reactions and incubated for 1 hour at room temperature with gentle rocking. Beads were then pelleted and washed 5 times with 1X Binding/Wash buffer, then resuspended in 1X Binding/Wash buffer.

On-bead protein digestion, LC/MS, and raw data analysis was done at the Stanford University Mass Spectrometry Core. Protein samples were digested on agarose beads. The proteins were reduced with 10 mM of dithriothreitol (DTT), incubated for five minutes at 55 °C, followed by head-over-head mixing at room temperature for 25 minutes using a Barnstead Thermolyne Labquake rotisserie shaker. They were then allowed to cool and alkylated with 30 mM acrylamide at room temperature for 30 minutes. This was followed by overnight digestion at 37 °C using 500 ng of mass spectrometry grade trypsin/LysC mix (Promega).

Post-digestion, samples were quenched with formic acid (adjusted to a pH ∼3) and desalted using MonoSpin C18 Solid-Phase Extraction (SPE) columns (GL Sciences). Finally, the samples were dried via SpeedVac (ThermoFisher Scientific) and exchanged into LC-MS reconstitution buffer (2% acetonitrile with 0.1% formic acid in water) for instrumental analysis.

Proteolytically digested peptides were separated using an in-house pulled and packed reversed phase analytical column (∼25 cm in length, 100 microns of I.D.), with Dr. Maisch 1.9-micron C18 beads as the stationary phase. Separation was performed with an 80-minute reverse-phase gradient (2-45% B, followed by a high-B wash) on an Acquity M-Class UPLC system (Waters Corporation, Milford, MA) at a flow rate of 300 nL/min. Mobile Phase A was 0.2% formic acid in water, while Mobile Phase B was 0.2% formic acid in acetonitrile. Ions were formed by electrospray ionization and analyzed by an Orbitrap Exploris 480 mass spectrometer (Thermo Scientific, San Jose, CA). The mass spectrometer was operated in a data-dependent mode using HCD fragmentation for MS/MS spectra generation.

The .RAW data were analyzed using Byonic v5.1.1 (Protein Metrics, Cupertino, CA) to identify peptides and infer proteins. A concatenated FASTA file containing Uniprot *Rattus norvegicus* proteins, bait sequences and other likely contaminants and impurities was used to generate an in-silico peptide library. Proteolysis with Trypsin/LysC was assumed to be fully specific. The precursor and fragment ion tolerances were both set to 12 ppm. Cysteine modified with propionamide was set as a fixed modification in the search. Variable modifications included oxidation on methionine and tryptophan, deamidation of glutamine and asparagine, and cyclization of glutamine and glutamic acid. Proteins were held to a false discovery rate of 1% using standard reverse-decoy technique (41). The final protein enrichment analysis of proteins bound to the MLS sequence vs the scrambled control sequence was carried out with the DESeq2 R package (42) using raw spectral counts as input.

### Conservation Analysis

We downloaded all available reference mammalian genomes (742 at time of writing) from the NCBI database and de novo annotated Mbp 3’ UTRs with sequence homology searching. First, we manually compiled 22 Mbp 3’ UTR sequences from representative species across mammalian clades to generate a seed multiple sequence alignment. We then used iterative nhmmer (43) searches with laxer E-value thresholds at each iteration (10^-10^, 10^-4^, then 10^-1^) to find 283 Mbp 3’ UTR homologues. To search for sub-sequences homologous to the Rattus norvegicus MLS amongst these homologous Mbp 3’ UTRs, we used iterative covariation-aware homology searching with the Infernal package (44). The first iteration was seeded with only the *R. norvegicus* MLS sequence and secondary structure. At each iteration, a homology model was built using the results from the prior iteration with the cmbuild program, calibrated with the cmcalibrate program, then used to search all remaining Mbp 3’ UTR homologues with the cmsearch program. The new hits were combined with hits from previous iterations and used to generate a new multiple sequence alignment with the CaCoFold algorithm (45). This new MSA was used as the input for the next iteration. This process was repeated until no new homologous MLS sequences were found. After the last iteration, 181 total MLS homologues were identified. We enforced a minimum E-value of 0.05 and only included hits on the sense strand. A final MSA was created using MAFFT Version 7 (46) and visualized with pyMSAviz (47).

## Supporting information

Dataset S01 (XLSX) Supporting Information Dataset S01_SLAP-seq

Dataset S02 (CSV) Supporting Information Dataset S02 Proteomics Raw Data

Dataset S03 (XLSX) Supporting Information Dataset S03_Species Sequence Alignment

Dataset S04 (XLXS) Supporting Information Dataset S04 _Synthesized Oligonucleotides

Table S1. Significantly enriched and de-enriched proteins in MLS pulldown study

## Acknowledgments

We thank members of the Barres, Fu, and Zuchero labs, including L. Meservey and G. Jones, for experimental guidance and scientific feedback; R. Rohatgi, C. Patel, G. Pusapati, and P. Harbury for experimental advice regarding RNA-protein pulldown and proteomics experiments; and I. Zheludev for insights into the execution and optimization of sequencing, sequencing analysis, and *in vitro* RNA-protein experiments. This work was supported by the National Institutes of Health (R35 GM122579 to R.D., R21 NS123533 to R.D. and J.B.Z., ZIA NS009432 to M.-M.F., P30 CA124435 for the Stanford Cancer Institute Proteomics/Mass Spectrometry Shared Resource), Human Frontier Science Program (RGEC32/2023 to M.-M.F.), National Multiple Sclerosis Society (postdoctoral fellowship to M.-M.F.), Dr. Miriam and Sheldon G. Adelson Medical Foundation (funding to Ben Barres), Myra Reinhard Foundation (funding to Ben Barres), and the Howard Hughes Medical Institute. Proteomics work was supported by the Vincent Coates Foundation Mass Spectrometry Laboratory, Stanford University Mass Spectrometry (RRID:SCR_017801) utilizing the Thermo Exploris 480 nanoLC/MS system (RRID:SCR_022215). This article is subject to HHMI’s Open Access to Publications policy. HHMI lab heads have previously granted a nonexclusive CC BY 4.0 license to the public and a sublicensable license to HHMI in their research articles. Pursuant to those licenses, the author-accepted manuscript of this article can be made freely available under a CC BY 4.0 license immediately upon publication.

