## Supplementary material for "RNA switch model for localization and translation of the myelin basic protein mRNA": Table S1. Significantly enriched and de-enriched proteins in MLS pulldown study

| Gene name | Protein name | $log_2$FoldChange | $p_{adj}$ |
|---|---|---|---|
| *Tars3* | Threonine-tRNA ligase | 1.59915872 | 0.00160976 |
| *Hnrnpf* | Heterogeneous nuclear ribonucleoprotein F | 1.47606397 | 0.00169003 |
| *Cct4* | T-complex protein 1 subunit delta | 1.11260304 | 0.00169003 |
| *Cct5* | T-complex protein 1 subunit epsilon | 1.10858645 | 0.00044553 |
| *Cct2* | T-complex protein 1 subunit beta | 1.09223648 | 0.00080849 |
| *Cct7* | T-complex protein 1 subunit eta | 1.07034113 | 0.00169003 |
| *Cct8* | T-complex protein 1 subunit theta | 0.9961152 | 0.00106049 |
| *Cct6a* | Chaperonin containing TCP1 subunit 6A | 0.94039895 | 0.0065329 |
| *Aimp1* | Aminoacyl tRNA synthetase complex-interacting multifunctional protein 1 | 0.92121638 | 0.02619575 |
| *Lars1* | Leucine-tRNA ligase | 0.85600888 | 0.007258 |
| *Cct3* | T-complex protein 1 subunit gamma | 0.8403121 | 0.01653504 |
| *Tcp1* | T-complex protein 1 subunit alpha | 0.68643992 | 0.06476284 |
| *Hadhb* | Trifunctional enzyme subunit beta, mitochondrial | −1.1105623 | 0.00478055 |
| *Igf2bp2* | Insulin-like growth factor 2 mRNA binding protein 2 | −1.4783708 | 0.009356 |
| *Abcf1* | ATP-binding cassette sub-family F member 1 | −1.5946004 | 0.00044553 |
| *Snrnp70* | U1 small nuclear ribonucleoprotein 70 kDa | −2.0235793 | 0.00809546 |
| *Msi2* | Musashi RNA-binding protein 2 | −2.165768 | 0.00654469 |
| *Sf3b2* | Splicing factor 3b, subunit 2 | −2.1681579 | 0.00545222 |
| *Sf3b3* | Splicing factor 3b, subunit 3 | −2.1719484 | 0.00106049 |

**Dataset S01 (XLSX) Supporting Information Dataset S01_SLAP-seq**
**Dataset S02 (CSV) Supporting Information Dataset S02_Proteomics Raw Data**
**Dataset S03 (XLSX) Supporting Information Dataset S03_Species Sequence Alignment**
**Dataset S04 (XLXS) Supporting Information Dataset S04_Synthesized Oligonucleotides**
